# Dynamic mechanisms of time-of-day-dependent adaptive immunity and vaccination responses

**DOI:** 10.1101/2025.05.19.654803

**Authors:** Xinyang Weng, Qi Ouyang, Hongli Wang

## Abstract

The timing of vaccine administration during the day significantly affects immunogenicity and efficacy, yet the mechanism governing the time-of-day dependent adaptive immunity and vaccine response remains elusive. Using mathematical modeling, we elucidate that the bistability arising from the self-enhancing homing process of antigen-presenting dendritic cells plays a key role in the time-of-day-dependent adaptive immune response. Modeling analyses of circadian-controlled immune responses to three vaccine types demonstrate that vaccine-specific differences in time-of-day-dependent immunity originate from distinctions in the circadian-regulated activation of antigen-presenting dendritic cells. This divergence is amplified by bistability in the dendritic cell homing process, resulting in long-term distinctions in adaptive immunity across vaccine types. The model results reveal a dynamic mechanism by which adaptive immune responses maintain circadian variations over extended time periods, suggesting that the timing of vaccine administration within the day is a promising strategy for effective disease prevention.

**Author Summary:** Infection with the same pathogen in the morning or afternoon, or receiving the same vaccine at different times of the day, often leads to significantly different consequences in terms of infection severity or vaccine efficacy. This phenomenon is known as the time-of-day-dependent immune response. Although it is relatively common, the underlying dynamic mechanisms remain mostly unclear. In this paper, we employ mathematical modeling to investigate in detail the dynamics of adaptive immune responses following vaccination. We reveal that the kinetic origin of the time-of-day-dependent immune response lies in the positive feedback process during the homing of antigen-presenting cells, particularly dendritic cells. The bistability induced by self-activation, under the regulation of the circadian clock, plays a decisive role in generating this diurnal time dependency. The discovery of this mechanism suggests that time-of-day-specific vaccination is an effective strategy to optimize vaccine efficacy.

## 1 Introduction

In mammals, the circadian clock regulates immunity through molecular interactions between its components and immune systems [1]. It influences various immunological processes [2,3], such as cytokine secretion [4,5], leukocyte migration [6,7], immune cell activation [8,9], and lymphocyte proliferation [10]. In experiments, both innate and adaptive immune responses were found to be highly dependent on the timing of initial exposure to antigens. The phenomenon of time-of-day-dependent immunity has been extensively observed in immune responses to lipopolysaccharide (LPS) stimulations [11–13], viral [14] and bacterial infections [15], as well as vaccinations [10,16,17].

Time-of-day differences in vaccination schedules can lead to long-term distinctions in adaptive immune response lasting for weeks or even months [16,18–20]. Morning vaccination of the influenza vaccine in humans, compared with afternoon vaccination, has been shown to elicit a higher antibody response one month after the initial challenge [20]. Subsequent research revealed similar time-of-day-dependent responses to vaccinations against bacillus Calmette-Guérin [19] and SARS-CoV-2 [16], with enhanced long-term immunity for when vaccinated in the morning. Mice vaccinated with human HAV vaccine and protein-based SARS-CoV-2 vaccine in the afternoon exhibited significantly increased frequency of specific T cells and antibody titers compared with those vaccinated at night [10]. Furthermore, the optimal timing for vaccination can be distinct for different vaccine delivery systems and adjuvants. It was reported that RNA-based SARS-CoV-2 vaccine elicited stronger antibody responses in humans when administered in the late afternoon [18]. The injection of bone marrow-derived dendritic cells (DCs) in mice triggered stronger adaptive immune responses at Zeitgeber Time 7 (ZT7) (daytime) [8,10], and another study showed that injection of ovalbumin (OVA) mixed with cytosine-phosphate-guanine (CpG) oligodeoxynucleotides (ODNs) in mice resulted in increased T cell activation and elevated levels of inflammatory cytokines at ZT19 (nighttime) [9].

Despite accumulating evidence of time-of-day-dependent immunity in vaccine administration, a quantitative understanding of the underlying mechanisms remains elusive. Unravelling the dynamic mechanisms is critical for people in modern society to adopt optimal time-of-day strategies in combating highly contagious and devastating diseases, such as the COVID-19 pandemic. Recently, the approach of mathematical modelling has been applied to investigate how the immune response depends dynamically on the time-of-day for initial challenge of viral infection [21]. It was demonstrated that the clock-controlled release of CCL2 and the incubation period of infection jointly determined the time-of-day-dependent innate immune response. In this paper, we construct a detailed model to investigate the dynamic mechanism underlying the time-of-day dependent immunity in vaccine administrations. The model incorporates the interactions between the circadian clock and the immune system known up to now, and considers three vaccine types—endotoxin-adjuvanted, nucleic-acid-adjuvanted, and DC-based vaccines—administered at different times of day. Our modeling successfully reproduces the experimentally observed data. Dynamic analyses demonstrate that the bistability in the circadian-controlled DC homing stage—a self-accelerating process—serves as the key mechanism governing time-of-day-dependent adaptive immune responses. Moreover, for different types of vaccines, the corresponding innate immune responses are regulated by the circadian clock in different ways. This difference is amplified by the bistability in the homing process of dendritic cells, thereby forming different time-of-day dependent immunity for different vaccine types. The model results present a dynamic mechanism for how the adaptive immune responses continue to exhibit circadian changes over long periods of time, which is significant in providing a strategy for using time-of-day to optimize vaccination efficacies.

## 2 Model

Vaccination provides long-term protection against pathogens by activating immune responses to produce antibodies and memory cells. A typical immune response, which is coupled with the circadian clock, includes the circulation of leukocytes in peripheral blood, the innate immune response in peripheral tissues, and the adaptive immune response within secondary lymphatic organs (such as draining lymph nodes, distant lymph nodes, and the spleen). The immune response exhibits variability depending on vaccine delivery systems and adjuvant types. In our modeling framework, we consider three categories: endotoxin-adjuvanted vaccines (*e.g*., antigen mixed with LPS), nucleic acid-adjuvanted vaccines (*e.g*., antigen mixed with CpG ODNs), and DC vaccines. In Fig 1, we illustrate the circadian clock-controlled interaction network between innate and adaptive immune responses for endotoxin-adjuvanted vaccines (see Fig S1 and Fig S2 in the Supporting Information for analogous networks of nucleic-acid-adjuvanted vaccines and DC vaccines, which differ primarily in the innate immune response module). The model is multicompartmental, encompassing peripheral tissues, secondary lymphatic organs, and peripheral blood (not included in Fig 1 for simplicity). The network comprises transcription factors, cytokines, and immune cells, along with their interactions in the processes of transcription and translation, cytokine secretion, cell proliferation and differentiation, and immune cell migration across compartments.

**Fig 1.**
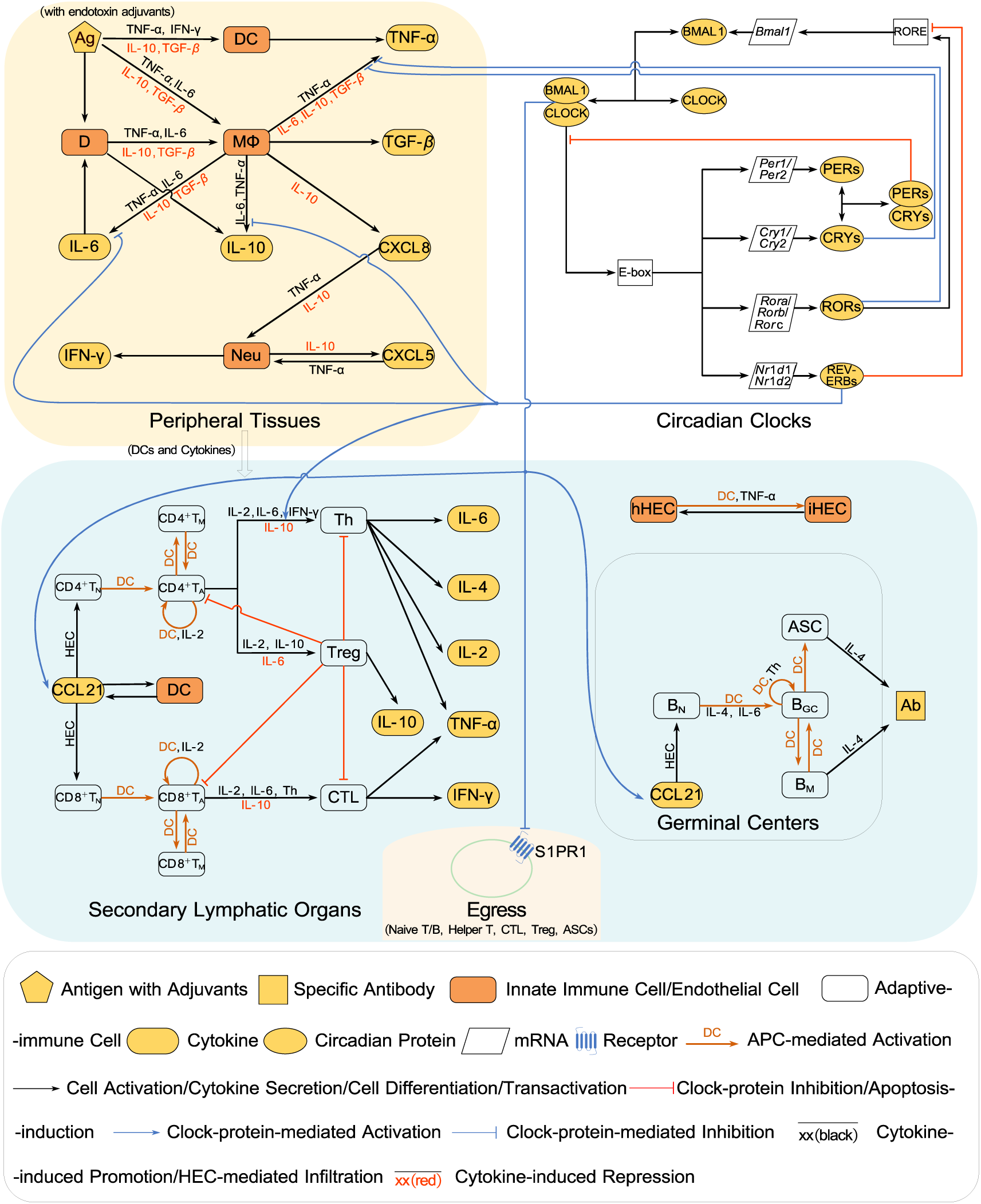
Schematic diagram depicting circadian-controlled immune response to endotoxin-**adjuvanted vaccines**. Gray region represents the circadian clock regulatory circuit. Yellow region indicates peripheral tissues at the vaccination site, corresponding to the primary innate immune activation zone. Blue region denotes secondary lymphoid organs, where adaptive immune responses (including humoral immunity in germinal centers) are initiated. The yellow region embedded in the blue depicts immune cell egress from secondary lymphatic organs into circulation. The antigen and immune adjuvants are collectively denoted with Ag. Homeostatic and inflammatory states of hematopoietic endothelial cells are designated as hHEC and iHEC, respectively. Subscripts N and M in adaptive immune cells denote naive lymphocytes (CD4^+^T_N_, CD8^+^T_N_ and B_N_) and memory cells (CD4^+^T_M_, CD8^+^T_M_ and B_M_), while subscripts A and GC indicate activated/proliferating lymphocytes (CD4^+^T_A_, CD8^+^T_A_) and B cells (B_GC_) in germinal center, respectively. Effector cells are abbreviated as: helper T cells (Th), cytotoxic T lymphocytes (CTLs), and antibody-secreting cells (ASCs). The peripheral blood functioning as a conduit for immune cell and cytokine trafficking between tissues and secondary lymphatic organs is omitted for simplicity.

In Fig 1, the innate immune response module resembles earlier models of acute immune response (see Ref. [22–24] but varies in its response to different vaccine types. For endotoxin-adjuvanted vaccines, dendritic cells and macrophages (MΦ in Fig 1) in the peripheral tissue at the vaccine injection site are activated, subsequently secreting substantial pro-inflammatory cytokines, such as tumor necrosis factor-alpha (TNF-α) and interleukin-6 (IL-6). These pro-inflammatory cytokines promote the activation of both macrophages and dendritic cells. Concurrently, macrophages secrete anti-inflammatory cytokines, such as IL-10 and transforming growth factor-β (TGF-β), which inhibit their own activation as well as that of dendritic cells, thereby reducing inflammatory cytokine secretion. Additionally, macrophages release chemokine CXCL8 to recruit neutrophils (Neu in Fig 1) for antigen clearance. The recruited neutrophils autonomously secrete CXCL5, establishing an amplification loop that enhances neutrophil recruitment and intensifies the inflammatory response. Peripheral tissue cells respond to antigenic signals and inflammatory mediators by causing tissue damage (D in Fig 1), which further stimulates macrophage activation. Activated DCs that have captured antigens migrate from peripheral tissues to draining lymph nodes through homing processes, thereby initiating the adaptive immune response. For nucleic-acid-adjuvanted vaccines in our modeling, the innate immune response is identical to that in Fig 1, except that the activation of DCs is extra repressed by the circadian clock as CLOCK-BMAL1 suppresses the expression of Toll like receptor 9 (TLR9) (see Fig S1 in the Supporting Information and Fig 2B) [9]. For DC vaccines, the pro-inflammatory effects of DC vaccines are not considered in our model (see Fig S2 in the Supporting Information). The DC cells, which have been loaded with antigens and activated, migrate straight to the lymph nodes and start the adaptive immune response. The module for the downstream adaptive immune response is common to the three vaccines considered in our model. The activated DCs are chemoattracted by CCL21 secreted by lymphatic endothelial cells (LECs) to enter afferent lymphatic vessels (LVs), in which DCs secrete chemokine CCL19 (not shown in Fig 1) and convert LECs-anchored CCL21 into soluble forms [25], thereby further enhancing DC homing. Upon reaching lymph nodes, DCs present antigens to naive lymphocytes (CD4^+^ T cells, CD8^+^ T cells, and B cells), initiating lymphocyte activation and proliferation. Most activated lymphocytes differentiate into effector cells (helper T cells, cytotoxic T lymphocytes, and antibody-secreting cells) that exit lymph nodes to execute immune functions, while the remainder develop into long-lived memory cells (memory CD4^+^ and CD8^+^ T cells, and memory B cells). Regulatory T cells suppress excessive immune responses, representing immune tolerance mechanisms. Moreover, DCs and the inflammatory cytokine TNF-α induce phenotypic transformation of hematopoietic endothelial cells (HECs) from homeostatic state to inflammatory state with enhanced permeability [26]. This transformation promotes lymphocyte infiltration from the peripheral blood into the lymph nodes and enhances the interaction between DCs and antigen-recognizing cells.

**Fig 2.**
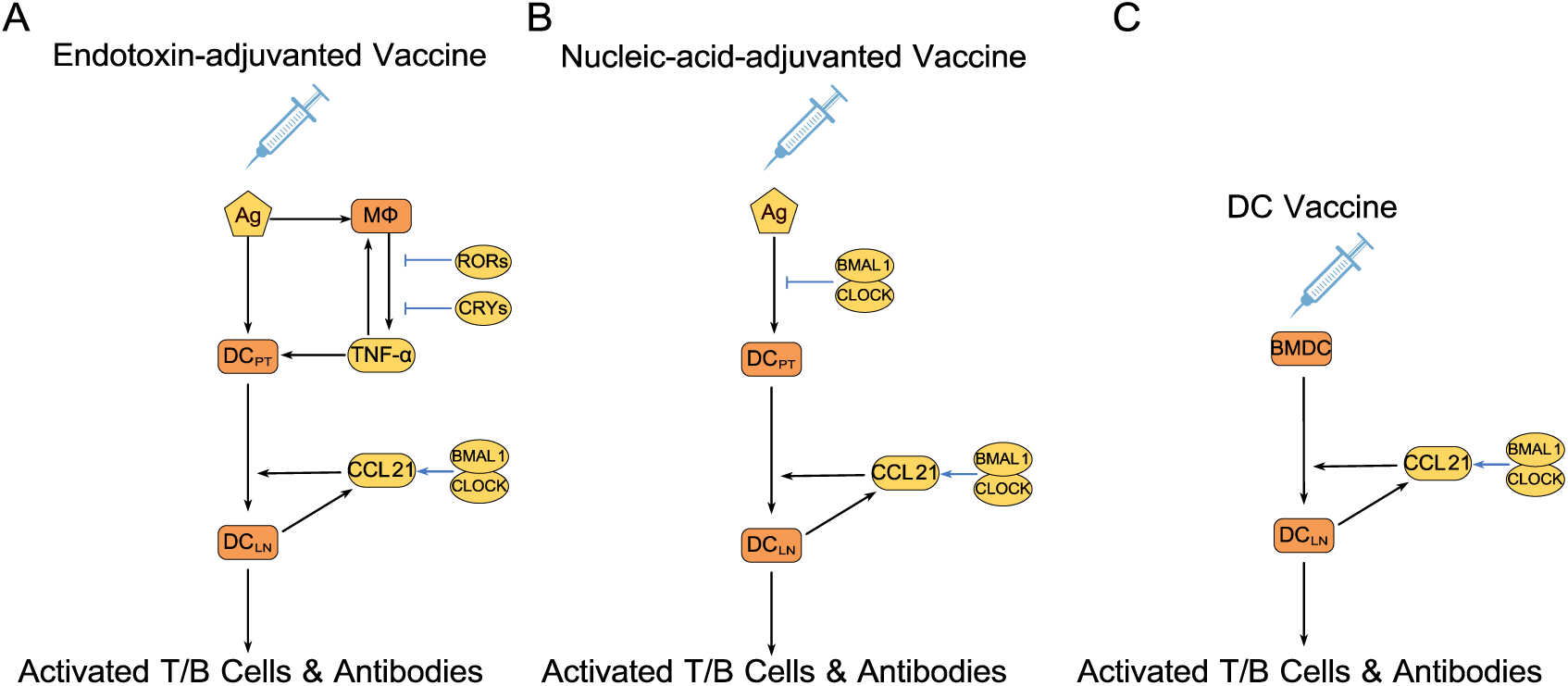
Circadian regulations of APC activation and homing that are pivotal for time-of-day-dependent adaptive immunity and vaccine responses. The key circuit governing APC activation and migration for: (A) endotoxin-adjuvanted vaccines, (B) nucleic-acid-adjuvanted vaccines, and (C) antigen-bearing dendritic cell vaccines. DC_PT_ indicates dendritic cells in peripheral tissues, while DC_LN_ represents dendritic cells in draining lymph nodes. The common feature of the circuits is that they all involve a CCL21-mediated self-accelerating DC homing process. The difference lies in the distinct regulations by the circadian clock during the activation of DCs.

In Fig 1, the module of circadian clock coupled with the immunity is primarily composed of two transcription-translation feedback loops that cooperatively interact to establish a self-sustained, robust 24-hour circadian rhythm (please refer to [27] for more detailed information). All immune cells in peripheral tissues and secondary lymphatic organs are assumed to share the same circadian clock. The regulatory interactions from the circadian clock to the immune system are denoted with blue arrows in Fig 1. In the innate immune response, the circadian clock component CRYs and RORs inhibit the expression of the pro-inflammatory cytokine TNF-α in macrophages [28–30]. REV-ERBs repress the transcription of inflammatory mediator IL-6 and anti-inflammatory cytokine IL10 [5,31]. For vaccines adjuvanted with nucleic acids, CLOCK-BMAL1 controls the activation of DCs as it suppresses the expression of TLR9 which is the receptor that mediates antigen-presenting cell (APC) activation. For endotoxin-adjuvanted vaccines, the receptor TLR4 that mediates DCs activation is not affected by the circadian clock [9,32]. In the adaptive immune response, CLOCK-BMAL1 promotes the expression of CCL21 in LECs to enhance DCs homing and lymphocyte migration [4]. It also downregulates the expression of egress receptor S1PR1, which is critical for lymphocyte trafficking [7]. REV-ERBα enhances the differentiation of T cells to Th17, a subset of T effector cells [33].

In our model, most immune cells and all the cytokines circulate in three compartments: peripheral tissues, secondary lymphoid organs, and peripheral blood (see Fig S3 in the Supporting Information). In a typical migratory pathway, immune cells infiltrate from blood vessels (not shown in Fig 1) into peripheral tissues, then home to draining lymph nodes via afferent lymphatic vessels, and exit through efferent lymphatic vessels into the blood [34]. In our model, the egress of activated lymphocytes (activated CD4^+^ T cells, CD8^+^ T cells, and B cells in germinal center) to the blood is inhibited due to the CD69-mediated repression of S1PR1, which is a critical receptor needed for lymphocyte trafficking. We assume that macrophages and neutrophils always stay in peripheral tissues for simplicity in our model.

## 3 Results

### 3.1 Dynamic simulations of time-of-day-dependent adaptive immunity and vaccination responses

#### 3.1.1 Modelling the steady-state immune system and innate immune response to the endotoxin challenge

We first simulated the dynamics of the immune system coupled with the circadian clock at steady state without the stimulation of antigens. Under homeostatic conditions, the ∼24-hour rhythmicity in the core components of the circadian clock regulates the expression of cytokines in immune cells, demonstrating a diurnal rhythm. The circadian rhythmic expression of chemokines leads to the diurnal rhythmic migration of lymphocytes between peripheral blood and lymph nodes [4,6,7]. The simulated circadian oscillations in the mRNA expression levels of the core circadian clock genes (*Bmal1*, *Cry*, *Ror*, *Rev-erbα*), CCL21 gene expression in lymph nodes, and the migration patterns of lymphocytes to lymph nodes are illustrated in Figs 3A, 3B, and 3C, respectively. The results are in good agreement with experimental observations in mice [7,35]. For *Bmal1*, *Rev-erbα*, *Ror*, and *Cry*, the peak mRNA levels occur at CT0, CT8, CT19, and CT20, respectively. The interlocking transcriptional feedback loops produce different phases of clock gene expression, leading to diverse patterns of clock-controlled immune functions. The CLOCK-BMAL1 complex generates the highest expression of CCL21 at CT9 (Fig 3B). CCL21, together with S1PR1, drives the circadian rhythmic migration of lymphocytes, which peaks at CT12 (Fig 3C). This agrees with experimental findings that lymphocyte homing to lymph nodes peaks at the onset of night [7].

**Fig 3.**
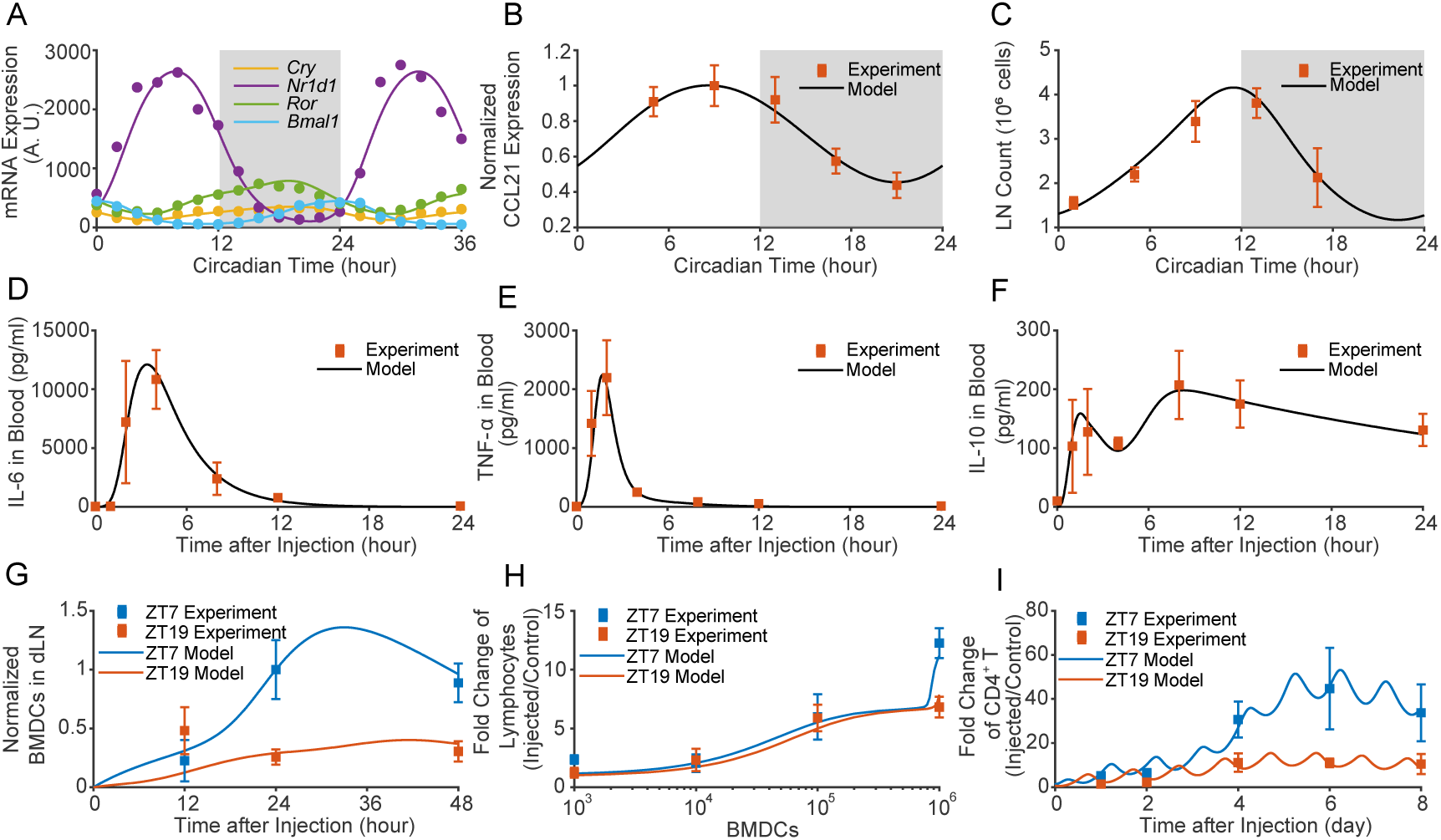
Simulation results of circadian rhythmicity in the circadian clock, immune cell trafficking, and the immune responses to endotoxin (LPS) stimulation and injection of bone-marrow-derived dendritic cell (BMDC). The curves represent simulation results, and the symbols denote experimental data measured in mice. (**A**) mRNA level oscillations in the expressions of four core circadian clock genes *Bmal1*, *Cry*, *Ror*, and *Rev-erbα*. (**B)** Circadian oscillations in CCL21 gene expression in lymph nodes. (**C**) Circadian rhythmicity in the homing dynamics of lymphocytes in lymph nodes. (**D-F**) Time-dependent changes in IL-6, TNF-α, and IL-10 levels in peripheral blood after LPS challenge. (**G**) Time evolutions of homed BMDCs in draining lymph nodes (dLNs) after subcutaneous injection of BMDCs. (**H**) Dose-dependent lymphocyte changes in dLNs at 48 h post-BMDC injection. (**I**) Time evolutions of CD4^+^ T cells in dLNs after BMDC injection. The experimental data: (A) from [35], (B,C) from [7], (D-F) from [24], and (G-I) from [10].

We next simulated the innate immune response coupled with the circadian clock in response to endotoxic LPS challenges. The acute immune response has been previously studied in rats with acute inflammation, in which varying doses of LPS endotoxin were intraperitoneally injected [24]. Since LPS primarily activates innate immunity, we excluded the adaptive immunity module in our modeling. The simulations reproduced the experimental results as depicted in Figs 3D-3F. The temporal changes in the concentrations of IL-6 (Fig 3D) and TNF-α (Fig 3E) exhibit pulsatile responses due to their rapid production and degradation, as well as the repression by the anti-inflammatory cytokine IL-10. In Fig 3F, the expression of IL-10 exhibits two distinct peaks, where the first peak is generated by the activation of macrophages, and the second peak is attributed to up-regulation mediated by damaged tissues. The dynamics of IL-10 could be altered due to the tight control of the circadian clock.

#### 3.1.2 Modelling the immune response to DC vaccines

The phenomenon of time-of-day-dependent adaptive immunity has been demonstrated in previous experiments of subcutaneous injection of BMDCs in mice [8,10]. The injection of antigen-bearing BMDC cells at ZT7 was found to induce a much stronger adaptive immune response compared to injection at ZT19. As shown in Figs 3G-3I, our simulations reproduced the experimentally observed time-of-day-dependent responses in adaptive immunity. The homing of BMDCs to draining lymph nodes was approximately threefold higher for injections at ZT7 (daytime) compared to ZT19 (nighttime) 48 hours post-injection (Fig 3G). The time-of-day dependence in infiltration of lymphocytes arose when the dosage of BMDCs exceeded 10^6^ counts (Fig 3H). Figs 3H and 3I depicted that the time-of-day-dependent homing dynamics of DCs further led to the infiltration of lymphocytes and activation of immune cells in a time-of-day-dependent manner. The number of activated CD4^+^ T cells was nearly three times higher for injections at ZT7. More detailed simulation results for the adaptive immune response to DC vaccinations are shown in Fig 4. The dynamic changes in CCL19 and CCL21 chemokine concentrations, DCs in draining lymph nodes, T helper cells, B cells in germinal centers, and antibody counts in peripheral blood all indicated a stronger immune response to vaccination at ZT7 than at ZT19. To measure the strength of the adaptive immune response, we adopted a score *ε*_*adaptive*_ based on immune cells in lymph nodes and cytokines in peripheral blood,

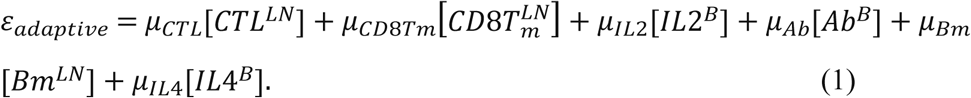

**Fig 4.**
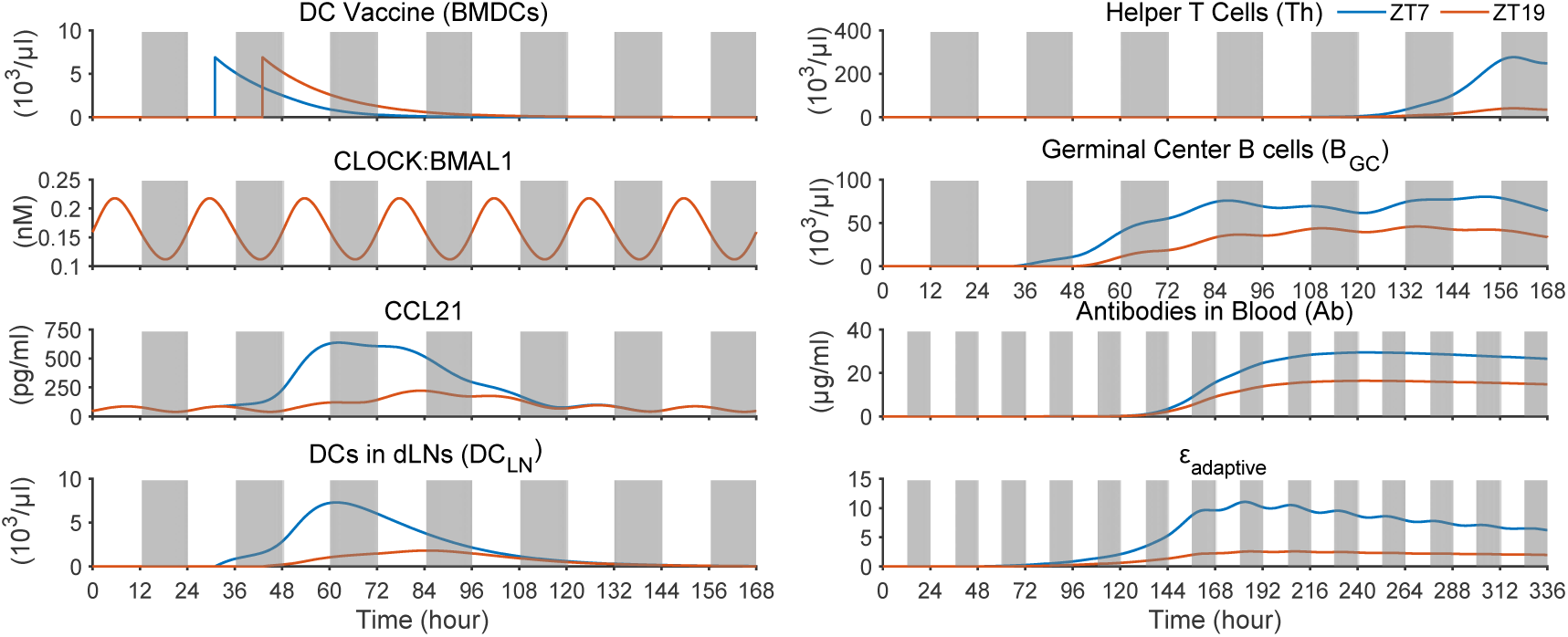
Simulated time evolutions demonstrate a time-of-day-dependent adaptive immune response to DC vaccines, with vaccinations administered at ZT7 eliciting a stronger immune response compared to those at ZT19.

As illustrated in Fig 4, the score *ε*_*adaptive*_ shows a prominent dependence on the vaccination time of day. From the simulated time evolutions in Fig 4, the differential immune response to vaccination timing originates from the differential behavior of chemokines: CLOCK-BMAL1, which peaks at ZT6 and troughs at ZT18, heavily shapes the circadian expression of CCL21, thereby promoting enhanced homing of DCs for vaccination at ZT7 compared to ZT19.

#### 3.1.3 Modeling the immune response to vaccines adjuvanted with endotoxin and nucleic acids

For endotoxin-adjuvanted vaccines, the endotoxins enhance the immunogenicity of vaccine antigens by activating the innate immune system and promoting antigen presentation. The immune response to endotoxin-adjuvanted vaccines was simulated for injections administered at ZT7 and ZT19, and the results are shown in Fig 5. Vaccination triggered a wave of macrophage activation, characterized by a pulse-like response. The secretion of TNF-α by macrophages was restricted by the intrinsic clock proteins CRYs (peaking at ZT21 and reaching a trough at ZT6) and RORs (peaking at ZT22 and reaching a trough at ZT10). For the case of ZT7 administration, the corresponding phase of inhibitors CRYs and RORs reaches their nadir, eliciting a greater amplitude of TNF-α release compared to ZT19. In Fig 5, the differential secretion of TNF-α was further transmitted to DC activation, and was markedly enhanced by the CLOCK-BMAL1-controlled DC homing dynamics. The difference persisted in subsequent lymphocyte recruitment and proliferation, ultimately leading to a long-term dependence of cellular immunity and antibody responses on the time-of-day of initial vaccine exposure. To evaluate inflammation during the immune response, we defined an inflammation score *ε*_*inflammation*_ based on signature immune cells in peripheral tissues and inflammatory cytokines in peripheral blood,

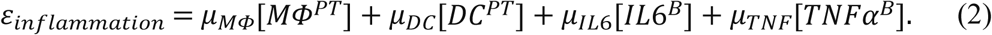

**Fig 5.**
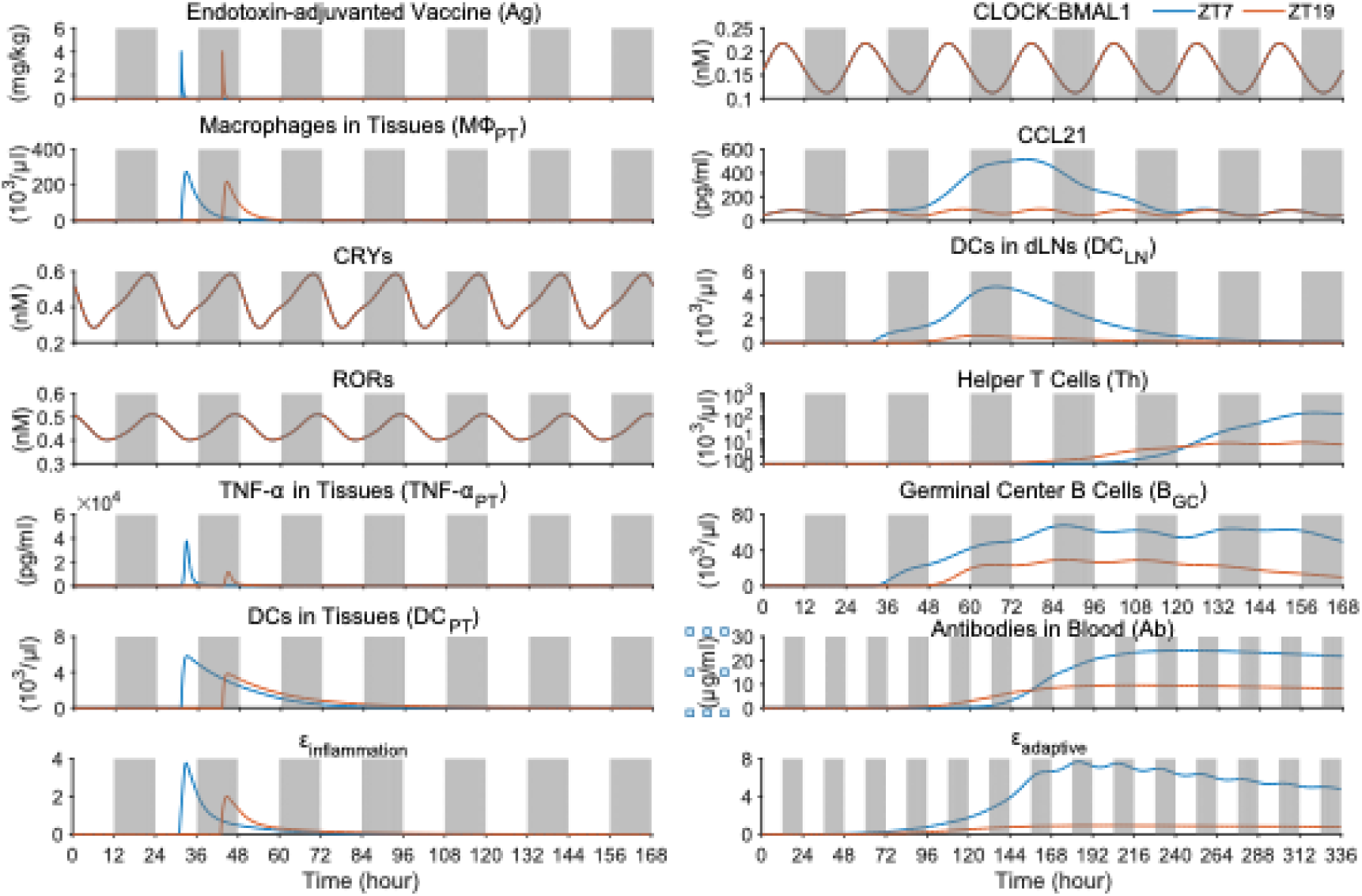
Simulated time evolutions demonstrate the time-of-day-dependent innate and adaptive immune responses to endotoxin-adjuvanted vaccines. Vaccinations administered at ZT7 elicit stronger immune responses compared to those at ZT19.

In Fig 5, both scores—*ε_inflammation_* (representing inflammation) and *ε*_*adaptive*_ (representing adaptive immune response)—demonstrated strong dependence on the time-of-day of the initial vaccine challenge.

The simulation results for immune response to nucleic-acid-adjuvanted vaccines are depicted in Fig 6. In this case, the activation of DCs as APCs is mediated by TLR9, whose transcription is regulated by CLOCK-BMAL1 [9,32] (see Fig 2 and Fig S1 in the Supporting Information). The simulated time evolutions are similar to those for endotoxin-adjuvanted vaccines, in which TNF-α and activated DCs in peripheral tissues, CCL21 cytokines and homed DCs in lymph nodes, B cells in germinal centers, antibody level in peripheral blood, and the collective indexes for inflammation and adaptive response all exhibit time-of-day dependence. The difference is that the time-of-day dependence is reversed in this case. In Fig 6, the immunity for vaccination at ZT19 is stronger than at ZT7, while in Fig 5, the stronger immunity occurs for the administration at ZT7. This difference should be attributed to the extra regulation from the circadian clock. The repression of CLOCK-BMAL1 on DC activation in the innate immune response is dominant in eliciting stronger inflammatory response and increased cellularity of DCs in peripheral tissues at ZT19 compared to ZT7. The resulted temporal difference in cellularity of tissue-endogenous DC overrides the clock-controlled effect on DC migration dynamics, resulting in reversed time-of-day-dependent patterns of adaptive immunity to vaccinations. Our simulation results indicate that immune responses elicited by different types of vaccines can exhibit distinct time-of-day dependencies due to the distinction in circadian clock regulation of the immunity response.

**Fig 6.**
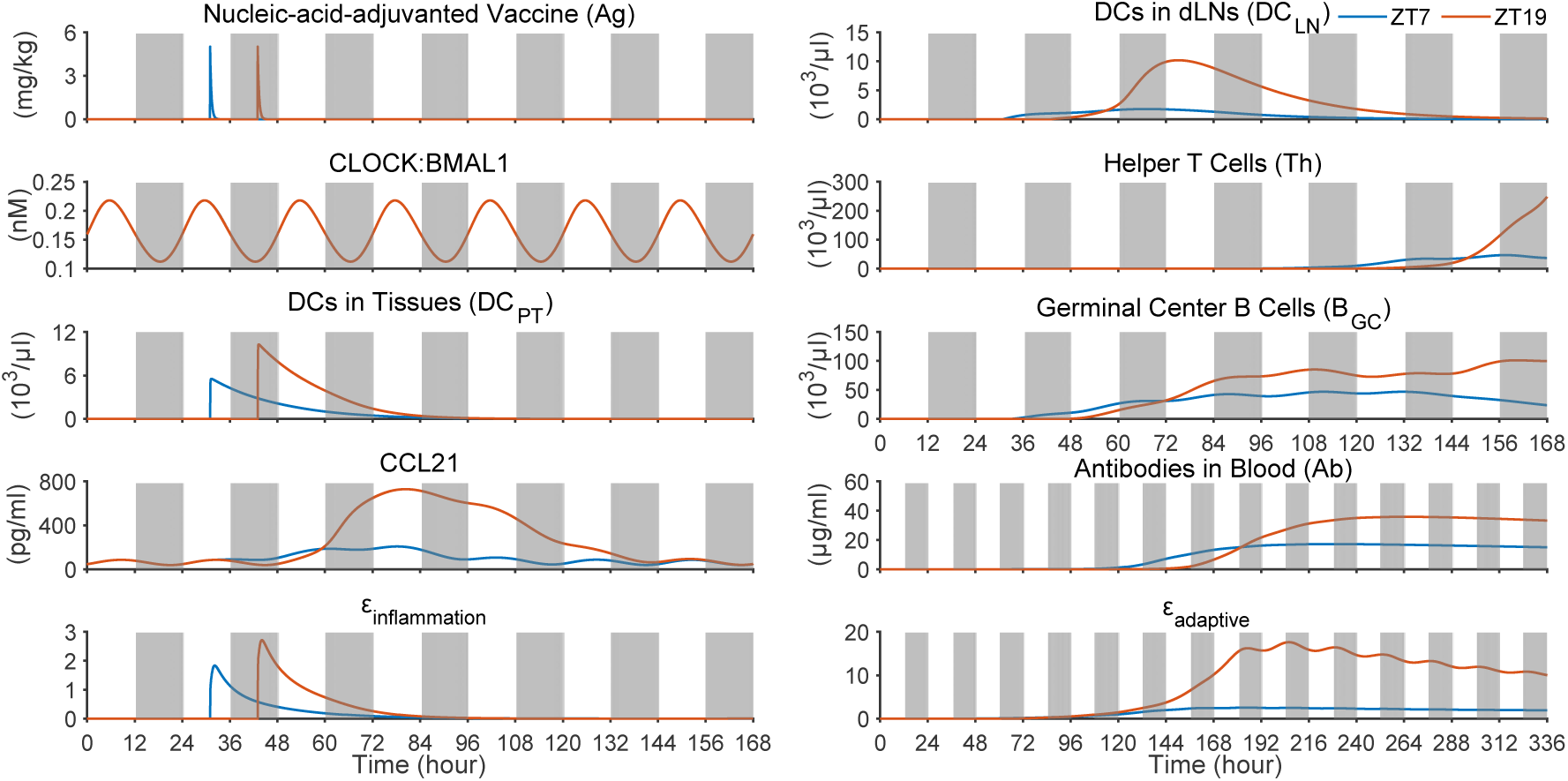
Simulations demonstrate that nucleic-acid-adjuvanted vaccines exhibit reversed time-of-day-dependent adaptive immunity compared to DC vaccines and endotoxin-adjuvanted vaccines, with stronger immune responses elicited at ZT19 versus ZT7.

### 3.2 Dynamic mechanisms of time-of-day dependent vaccine immunity

The time-of-day dependence of immune responses to vaccinations simulated above is rooted in the circadian regulation of immunity, mediated by clock proteins such as CLOCK-BMAL1, RORs, CRYs, and REV-ERBs. The activation and migration of antigen-presenting cells play a key role in triggering adaptive immune responses. The circadian control of dendritic cell activation and homing is central to time-of-day-dependent immunity. Fig 2 depicts the major circadian regulations governing DC activation and homing process across three vaccine types. They have distinct characteristics in DC activation processes. For endotoxin-adjuvanted vaccines, DC activation is modulated by RORs and CRYs, while for nucleic-acid-adjuvanted vaccines, it is controlled by CLOCK-BMAL1. Pre-activated DCs bypass the circadian regulation for DC vaccines. A self-accelerating mechanism mediated by chemokine CCL21 drives DC homing from peripheral tissues to draining lymph nodes, a process shared by all vaccines. Critically, this positive feedback loop is circadian-regulated via CLOCK-BMAL1, which promotes CCL21 expression in lymphatic endothelial cells, thereby enhancing DC migration. The resulting bistability (from self-accelerated dynamics) underpins the time-of-day-dependent adaptive immunity.

The mechanism underlying the time-of-day-dependent vaccination response can be elucidated through dynamic analyses. In the positive feedback loop of DC homing, activated DCs—recruited via CCL21 chemotaxis—secrete chemokine CCL19 (not included explicitly as a dynamic variable in our model), which converts LEC-anchored CCL21 into soluble forms, thereby further enhancing DC homing. The dynamics of DC homing process are governed by the following system of nonlinear ordinary differential equations (ODEs),

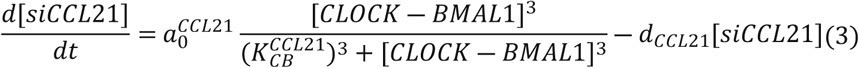

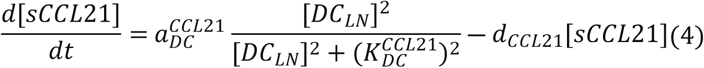

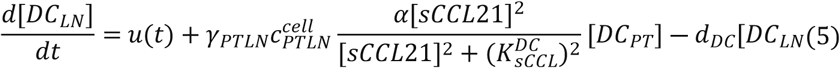

where

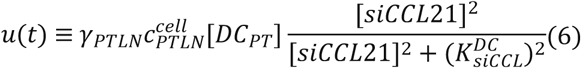

In Eqs. 3 and 4, [*siCCL21*] and [*sCCL21*] represent the concentrations of chemokine CCL21 in its surface-anchored form on LECs and soluble form, respectively. In Eq. 5, [*DC_LN_*] represents the number of DCs that have homed to draining lymph nodes, while [*DC_PT_*] represents DCs in peripheral tissues, which exhibit temporal decay (see Fig 6). *u*(*t*) in Eq. 5 is defined in Eq. 6, with the effects of [*DC_LN_*] and [*siCCL21*] incorporated. The self-accelerated DC homing process is described by Eqs. 4 and 5, which together form a nonautonomous system driven by the time-dependent variables *u*(*t*) and [*DC_PT_*](*t*). To analyze the dynamics governed by these equations, *u* and [*DC_PT_*] are treated as control parameters, and a bifurcation diagram is constructed in the *u*-[*DC_PT_*] parameter plane (Fig 7A). The surface of steady states in the bifurcation diagram reveals a bistable regime bounded by saddle-node bifurcation curves, which originate from a cusp bifurcation (approximately at (*u*, [*DC_PT_*]) = (1.32,0.28)). This bistability is further elucidated in one-parameter bifurcation diagrams (Figs 7C and 7D), generated by taking cross-sections of the two-dimensional bifurcation diagram. Also shown in Fig 7A are the time evolution trajectories for DC vaccines administered at ZT7 and ZT19, along with their projections onto the *u*-[*DC_PT_*] plane, as depicted in Fig 7B.

**Fig 7.**
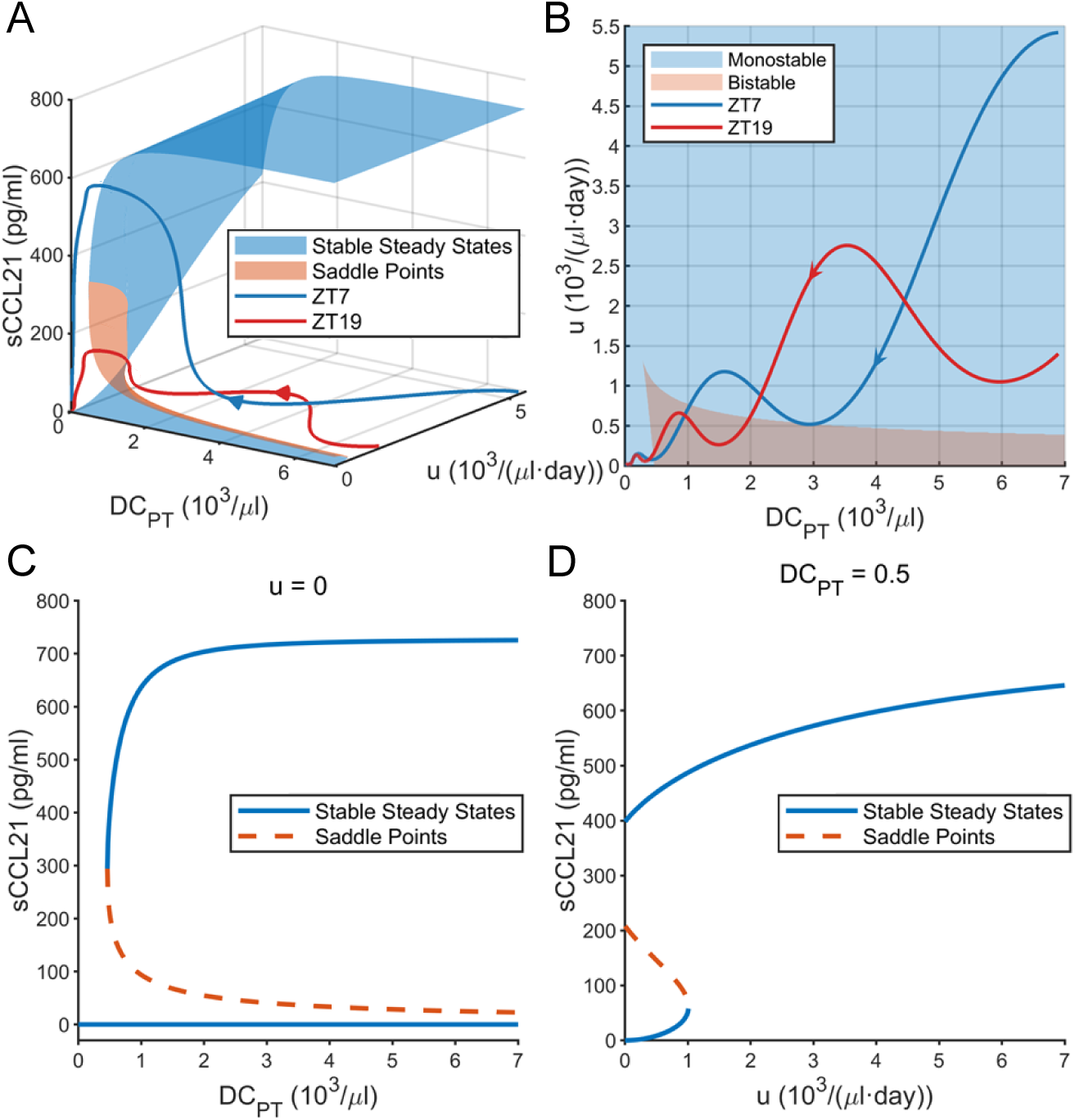
Dynamic bifurcation diagram for the isolated subsystem of Eqs. 4 and 5 for the self-accelerating DC homing process. (A) Bifurcation diagram in the *u*-[*DC_PT_*] plane demonstrating bistability, along with the two trajectories for DC vaccines administered at ZT7 and ZT19. (B) Bifurcation diagram and the trajectories projected onto the *u*-[*DC_PT_*] plane. (C) Saddle-node bifurcation diagram as [*DC_PT_*] is varied with *u* fixed at 0. (D) Bifurcation diagram as *u* is controlled with fixed [DC_PT_]= 0.5 (10^3^/*μl*). The arrows in (A) and (B) indicate the direction of time evolution.

As shown in Fig 7B, the time-dependent variables *u*(*t*) and [*DC_PT_*](*t*) transiently enter the bistable regime before decaying to zero. The time-of-day-dependent adaptive immune response to vaccinations stems from these transient bistable states, which are governed by *u*(*t*)—a parameter strongly modulated by the circadian oscillations of the CLOCK-BMAL1 protein complex—and the monotonically decaying [*DC_PT_*]. Fig 8 shows the phase-plane trajectories of [*DC*_*LN*_] and [*sCCL*21] during four sequential phases of the adaptive immune response to DC vaccines administered at ZT7 (blue) and ZT19 (red). In the early stages (Figs 8A, B), the bistable regime—comprising low and high steady states (black dots)—emerges from the intersection of nullclines (dashed lines) for *d*[*DC*_*LN*_]/*dt* = 0 and *d*[*sCCL*2]/*dt* = 0. The trajectories start from the low steady state upon the activation of DC homing. Once trajectories (blue line for ZT7) cross the saddle point separating the low and high steady states, they are driven by the saddle’s unstable manifold (gray line) toward the high steady state. This causes them to diverge from the trajectories (red line for ZT19) that fall in the basin of attraction of the low stable steady state without crossing the separatrix between the basins of attraction for the two stable steady states. As [*DC_PT_*] decays during the process of DC homing, the two nullclines separate at their high ends, causing the high steady state and the saddle to disappear through a reverse saddle-node bifurcation (see Figs 8C and 8D; see also Figs 7C and 7D). Consequently, all trajectories return to the low stable steady state that persists indefinitely. For a specific vaccine, the administration time of day determines whether the trajectory of the DC homing process is transiently driven by the saddle’s unstable manifold toward the high stable steady states. An optimal time-of-day strategy exists for each vaccine to maximize the vaccine efficacy. This represents the dynamic mechanism underlying the phenomenon of time-of-day-dependent development and maintenance of adaptive immune response to vaccinations.

**Fig 8.**
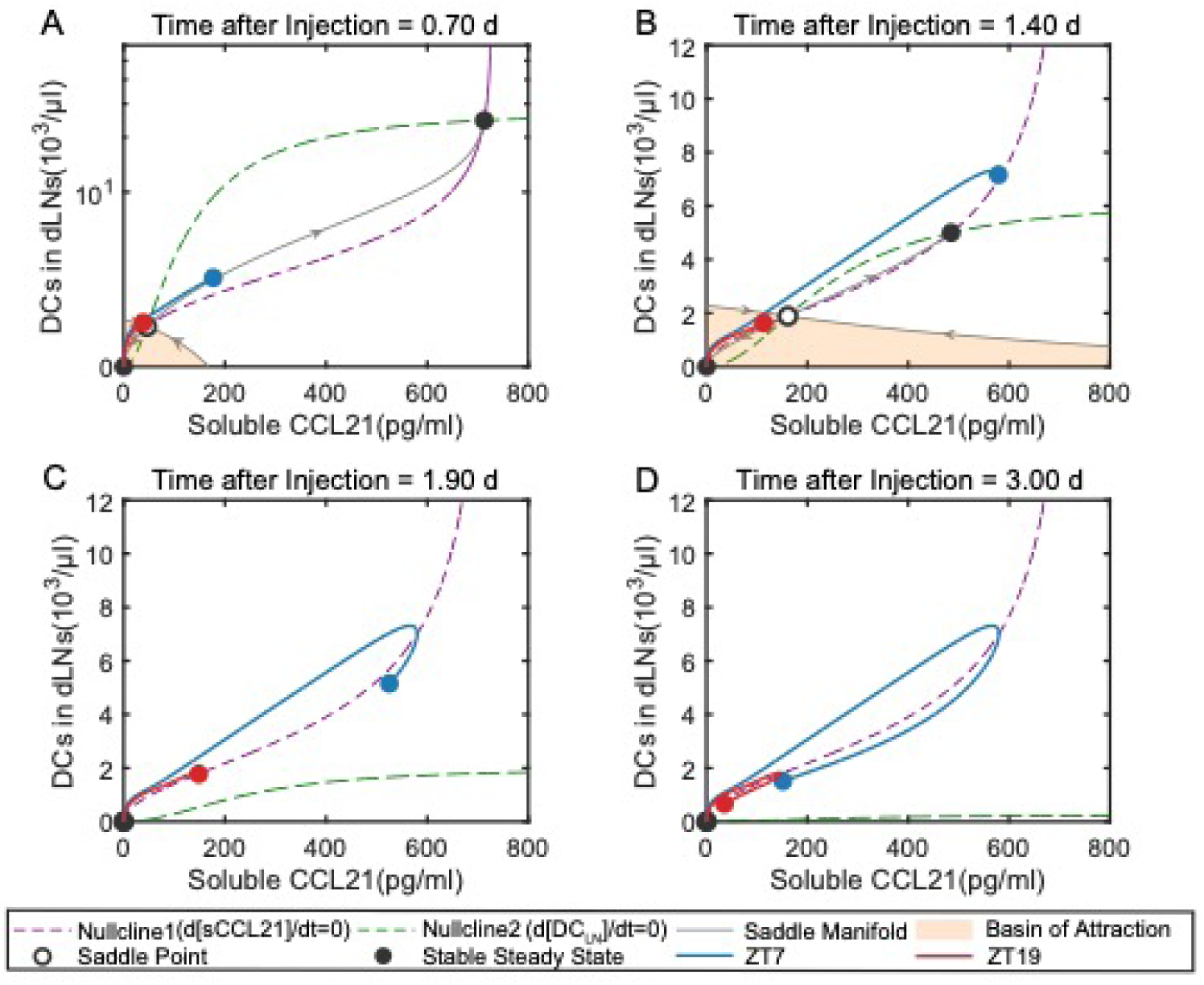
Simulated DC homing process (described by Eqs. 4-5) reveals a much stronger response following DC vaccine administration at ZT7 versus ZT19.

For different vaccines, the time-of-day dependence varies because the time evolution of DC homing also depends on the ways in which the APC activation is regulated by the circadian clock. Fig 9 depicts the trajectories of ([DC_LN_], [sCCL21]) of DC homing for nucleic-acid-adjuvanted vaccine administered at ZT7 and ZT19. The response magnitudes are exchanged between the two cases, as the administration at ZT19 enters the attractive basin of the high steady state instead. The difference stems from the distinct circadian regulations. For DC vaccines, pre-activated DCs migrate directly to lymph nodes without circadian control (see Fig 2C), whereas for nucleic-acid-adjuvanted vaccines, DC activation is regulated by CLOCK-BMAL1 before homing to lymph nodes (see Fig 2B). In Fig 6, the activation of DCs in peripheral tissues is less repressed for the vaccine administered at ZT19, giving it an advantage over ZT7. This advantage is further enhanced by the mechanism of transient bistability, as illustrated in Fig 9. The origin of time-of-day dependent adaptive immune response to endotoxin-adjuvanted vaccines is similar to the case of DC vaccines, as also depicted in Fig S4 in the Supporting Information. The result that circadian-regulated APC activation and migration are pivotal for time-of-day-dependent adaptive immunity is in agreement with the sensitivity analyses, in which the parameters related to the circadian clock-controlled dendritic cell activation 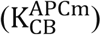 and homing 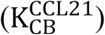 have thegreatest impact on the adaptive response (see Fig S4 in the Supporting Information).

**Fig 9.**
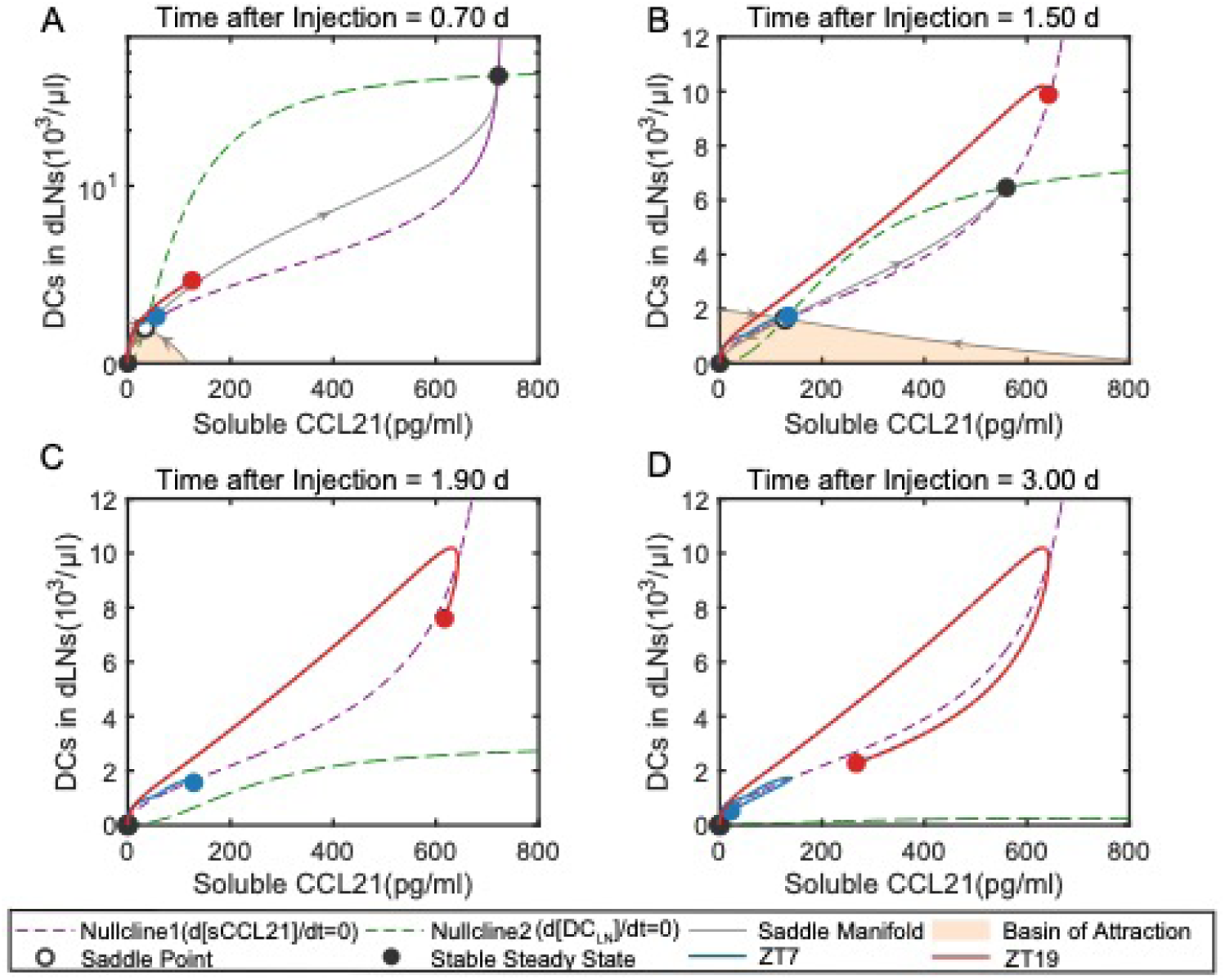
Transient dynamic behaviors demonstrate time-of-day-dependent dendritic cell homing in response to nucleic-acid-adjuvanted vaccines. The results exhibit a reversed situation compared to Fig 8, with a much stronger response following administration at ZT19 versus ZT7.

## 4 Discussion

The mechanisms behind the time-of-day-dependent adaptive immune responses to vaccination are fascinating. It is particularly intriguing to understand how these circadian differences can remain intact even weeks after the initial challenge. We have developed a comprehensive model to explore the dynamic mechanisms for why adaptive immune responses continue to exhibit circadian changes over long periods of time. The model integrated the current knowledges on the interplay between the circadian clock and the immune response, and examined different vaccine types. Our simulations reproduced the experimentally observed results of circadian rhythmicity under homeostatic conditions and long-term, time-of-day-dependent adaptive immune responses. By checking into the circadian-regulated immunological dynamics, including innate immune activation, adaptive lymphocyte trafficking, T cell activation/differentiation, and B cell-mediated antibody production, we demonstrated that clock-regulated dendritic cell activation and homing represent the most critical steps in time-of-day dependent adaptive immunity. Dynamic analyses highlighted that the circadian control of DC homing, which is mediated by the clock protein CLOCK-BMAL1 that enhances the expression of chemokine CCL21 for DC migration, is a self-accelerating process that generates bistability in its dynamics. This enables bistable immune states of low and high levels of adaptive immunity. Such bistable dynamics explains the dose-dependent divergence in lymphocyte infiltrations observed in experiments [10], and is in line with the previous finding that circadian disruption in DC-specific *Bmal1* knockout mice abolishes rhythmic T cell activation and antibody production [8, 36].

Our simulations of three vaccine types—endotoxin-adjuvanted, nucleic acid-adjuvanted, and dendritic-cell-based vaccines—revealed that the time of day differentially impacts adaptive immune responses across vaccine types. This is in agreement with previous findings that the optimal time of day of vaccination is dependent on various factors, including the type of vaccine, the immune adjuvants adopted, and specific immunization protocols. It was reported that immunization with ovalbumin (OVA)-loaded DCs [8], myelin oligodendrocyte glycoprotein (MOG) with Complete Freund’s Adjuvant (CFA) [7] or OVA adjuvanted with whole-cell pertussis (wcP) [37] in mice induced stronger T-lymphocyte activation when administered during daytime phases (ZT6-ZT8), while injection of OVA-CpG [9], OVA-Alum [38] or NP-conjugated to chicken gamma globulin (NP-CGG) with monophosphoryl lipid A (*i.e.*, MPL which is a TLR4 agonist) as an adjuvant [39] during the nighttime (ZT16-ZT19) elicited increased frequency of activated lymphocytes and higher antibody responses. Our model suggests that the difference in the optimal time of day for vaccinations arises from circadian regulation in both innate and adaptive immune responses. At the stage of innate immune response, the strength of APC activation relies on the phase of the circadian clock and therefore depends on the time of day of vaccination. The time-of-day-dependent variation in APC activation is further amplified by the bistability of the DC-homing positive feedback loop. This ultimately leads to sustained differences in antigen-specific T and B cell responses depending on the time of day of vaccination. Different vaccine types typically exhibit distinct circadian regulation patterns in antigen-presenting cell activation, as shown for the three vaccine types in Fig 2. For DC-based vaccines, APCs are pre-activated, and the time-of-day dependence in adaptive immune responses directly results from CLOCK-BMAL1-mediated circadian regulation of bistability. For endotoxin-adjuvanted vaccines, lower CRY/ROR activity at the trough phase enhances the TNF-α-driven DC activation, whereas nucleic-acid-based vaccines exhibit stronger nighttime responses due to the CLOCK-BMAL1 repression of TLR9 signaling. The variations lead to different time-of-day-dependent APC activation patterns, thereby contributing to the difference in time-of-day-dependent vaccination efficacy for different vaccines.

In summary, we elucidated by using the approach of mathematical modelling that the time-of-day dependent vaccination efficacy is rooted ultimately in the bistability in APC homing dynamics. The bistability, combined with the time of day of vaccination and the circadian regulation of APC activation, collectively contributes to the time-of-day-dependent vaccine efficacy. This provided a theoretical framework for understanding how short-time period circadian cues propagate into the long-term adaptive immune memory and why temporal variations in antibody titers or memory B cells persist weeks post the initial vaccination [10, 16]. Although there are plenty of factors that influence humoral and cellular vaccine responses in humans, such as vaccine factors (vaccine type, adjuvant, and dose) and administration factors (schedule, site, route, time of vaccination, and co-administered vaccines and other drugs). The model results presented here highlighted that the timing of vaccination administration offers an effective way to improve vaccine immunogenicity and efficacy.

## 5 Method

The model consists of three compartments: peripheral tissues, secondary lymphoid organs, and blood vessels (see Fig S3 in the Supporting Information). The dynamics of the circadian-controlled immune response to vaccinations are described by a system of coupled nonlinear ODEs (see Eqs. S1–S4 in the Supporting Information). The dynamic equations mathematically represent multiple immunological dynamics, encompassing processes such as immune cell activation, cytokine secretion, cell migration, cytokine diffusion, cell proliferation, molecular degradation, and transcriptional regulation. Hill functions are extensively employed. Take Eq. S2.3 for an instance, the activation of naive CD4^+^ T cells by DCs in lymph nodes is described by the form ∼[*DC*]/([*DC*]+ K) ·[naive CD4T] multiplied by ^[IL2^LN^]^/^(*K*+ [IL2^LN^])^, as it is IL-2 facilitated. The IL-2 facilitated cell proliferation rate takes the form of Logistic model ∼[*CD*4*T*] (1 ―[CD4T]/*N*) multiplied by ∼^[IL2^LN^]^/^(*K*+ [IL2^LN^])^·^[*DC*]^/^([*DC*]+ *K*)^. The rate of cytokine secretion by DCs is represented as ∼[*DC*]^2^ /([*DC*]^2^+ ^K2^), as shown in Eq. S2.18. For cell migration between compartments, we take Eq. S2.2 as an example. The movement of naive CD4^+^T cell from the blood to the secondary lymphatic organs is guided by the chemokine CCL21 and depends on the permeability of the HEV endothelial cells (HEVECs). The influx takes the form ∼γ[HEC]·α[CCL21_LN_]^2^/(*K*^2^ + [CCL21]^2^)·[naive CD4T]. In Eq. S2.2, the migration of naive CD ^+^T cell from the secondary lymphatic organs to the blood is mediated by S1PR1 receptor on the surface of lymphocytes, and the efflux takes the form ∼[*S*1*PR*1]^3^/([*S*1*PR*1]^3^ + *K*^3^)· [naive CD4T]. Due to the constraint of mass conservation, the influx of cells and cytokines migrating from the blood to the secondary lymphatic organs is scaled with factor *γ*_*BLN*_. It is estimated as the volume ratio of the peripheral blood (1700μl) to a single lymphatic node (∼8μl) for mouse. The homing of dendritic cells from peripheral tissues to secondary lymphoid organs is modeled by Eqs. 3-6 (i.e., Eqs. S2.1, S2.17, and S2.18 in the Supporting Information). To model the circadian clock, we adapted the mathematical framework proposed by Abo et al. [22], extending it by incorporating additional coupling terms to link circadian rhythms with immunological dynamics. The complete model comprises 75 state variables and 288 parameters. We employed the MATLAB *ode45* solver to numerically integrate the ODEs. The complete set of equations and parameter values are detailed in the Supporting Information.

In our model, a subset of parameters is experimentally based or adopted from existing models in the literature, with the remaining parameters being estimated by fitting to experimental data. The fitted parameters were calibrated through a four-step procedure. First, the circadian clock module was decoupled from the immune system dynamics. Parameters adopted in the mathematical model of the circadian clock by Abo et al. [22] were implemented to establish autonomous circadian rhythmicity (see Fig 3A, and Eqs S4.1-12 in the Supporting Information for the ODEs). Next, dynamic fitting was performed to precisely reproduce the experimental data [7] for circadian rhythmicity in lymphocyte homing and chemokine expression in lymph nodes without antigen stimulation (see Figs 3B and 3C and Eqs. S2.2, S2.5, S2.9, S2.12, S2.17, S2.19, S3.2-S3.4, S4.1-12 in the Supporting Information). At this stage (and here after), the regulation of circadian clock was turned on, and a subset of parameters associated with innate immunity were determined based on experimental data. The remaining parameters of the innate immune response were calibrated using cytokine data (IL-6, TNF-α, IL-10) from LPS-challenged rat experiments (see Figs 3D-F) [24]. During this step, adaptive immunity was excluded from the model by fixing all kinetic parameters associated with adaptive-innate immunity interactions to zero during data fitting. Finally, the full model was validated by using DC vaccination data in mice [10] to fit the time-of-day-dependent adaptive responses to resolve the undetermined parameters (see Figs 3G-I). All numerical optimizations were done by utilizing MATLAB *fminsearch* function. With these determined parameters, the results in Fig 4, Fig, 5 and Fig 6 were obtained by simulating ODEs of DC vaccine responses (Eqs. S1.2, S1.5-S1.8, S1.11-S1.17, S2.1-S2.21, S2.23-S2.27, S3.2-S3.14 and S4.1-S4.12), endotoxin-adjuvanted vaccine response (Eqs. S1-S4), and nucleic-acid-adjuvanted vaccine response (Eqs. S1-S4) in the Supporting Information, respectively.

The scores *ε*_*adaptive*_ and *ε*_*inflammation*_ defined in Eq. 1 and Eq. 2 were employed to systematically evaluate the circadian clock’s regulatory effects on immunity, where *ε*_*adaptive*_ represents the adaptive immune response strength and *ε*_*inflammation*_ corresponds to the inflammatory activity level. In Eq. 1, the concentrations of antibody [*Ab*^*B*^] and B memory cells [*Bm*^*LN*^] represent the humoral immunity level of the immune system; cytotoxic T lymphocytes [*CTL*^*LN*^] and CD8⁺ memory T cells [*CD*8*T*^*LN*^] represent the cellular immunity level; and cytokines related to the adaptive immune response [*IL*2^*B*^] and [*IL*4^*B*^] represent the activation degree of the adaptive immune system. The superscripts “PT,” “LN,” and “B” denote peripheral tissue, lymphoid tissues/organs, and peripheral blood, respectively. In Eq. 2, the macrophage population [*Mφ*^*PT*^] and inflammatory cytokines [*IL*6^*B*^] and [*TNFα*^*B*^] quantify the inflammatory activity of the immune system, while dendritic cells [*DC*^*PT*^] characterize its antigen-presenting capacity. In Eqs. 1-2, the weighting coefficient μ was fixed as the reciprocal of the peak values of the corresponding immune components after the immune system responded to endotoxin-adjuvanted vaccines (at a dose of 3 mg/kg). The numerical values of the weighting coefficient μ are provided in the Supplementary Information.

## Funding

This work was supported by the National Natural Science Foundation of China (12090051 to HW), and the Starry Night Science Fund of Zhejiang University Shanghai Institute for Advanced Study (to QO). The funders had no role in study design, data collection and analysis, decision to publish, or preparation of the manuscript.

## Data availability

The source codes used to produce the results and analyses in this manuscript are available on GitHub at https://github.com/npw913/Circa_immune_simulation.

## Author Contributions

**Conceptualization**: Hongli Wang, Qi Ouyang.

**Formal analysis**: Xinyang Weng.

**Funding acquisition**: Qi Ouyang, Hongli Wang.

**Investigation**: Xinyang Weng.

**Methodology**: Xinyang Weng, Hongli Wang, Qi Ouyang.

**Software**: Xinyang Weng.

**Supervision**: Hongli Wang, Qi Ouyang

**Validation**: Xinyang Weng, Hongli Wang.

**Visualization**: Xinyang Weng.

**Writing – original draft**: Xinyang Weng.

**Writing – review & editing**: Hongli Wang, Xinyang Weng.

## Supporting information

**Text S1.** Supporting information, including Figures A-E, Equations A-D, and Tables A-J.

## Notes

### Competing Interest Statement

The authors have declared no competing interest.

